# The ABI3-ERF1 module mediates ABA-auxin crosstalk to regulate lateral root emergence

**DOI:** 10.1101/2023.03.08.531793

**Authors:** Jing Zhang, Pingxia Zhao, Siyan Chen, Liangqi Sun, Jieli Mao, Shutang Tan, Chengbin Xiang

**Author notes:** Corresponding authors: Pingxia Zhao, Chengbin Xiang. Jing Zhang and Pingxia Zhao contributed equally.

## Abstract

Lateral root (LR) development is regulated by hormones and environmental signals. Abscisic acid (ABA) is involved in LR development, but it is not well understood how ABA signaling interacts with auxin to regulate LR formation. Here, we report that *Arabidopsis ERF1* is responsive to ABA during LR development and mediates the crosstalk between ABA and auxin signaling to regulate LR emergence. ABA attenuates ERF1-mediated excessive auxin accumulation with altered distribution known to inhibit LR emergence. ABI3 acts as a negative factor in LR emergence and transcriptionally activates *ERF1* by binding to the RY motif in its promoter, and reciprocally, ERF1 activates *ABI3*, which forms a positive regulatory loop that enables rapid signal amplification. Notably, ABI3 physically interacts with ERF1, consequently reducing the *cis* element-binding activities of both ERF1 and ABI3 and thus attenuating ERF1-regulated transcription of *PIN1*, *AUX1*, and *ARF7* involved in the modulation of LR emergence and ABI3-regulated *ABI5* in ABA signaling, which may provide a molecular rheostat to avoid overamplification of auxin and ABA signaling. Taken together, our findings unveil the pivotal ABI3-ERF1 module that mediates the crosstalk between ABA and auxin signaling in LR emergence.

## Introduction

Root systems display variable architectures that contribute to survival strategies of plants. Lateral roots (LRs), as branched root systems, are major architectural determinants of water and nutrient uptake from soil.^1–3^ To date, LR development has been well characterized in *Arabidopsis thaliana*. LRs branch off the primary root, initiating from pericycle founder cells located at xylem poles, forming a dome-shaped primordium, eventually breaking through the epidermis and generating a new functional root.^4–6^

Auxin is essential for LR initiation and emergence.^7–9^ Auxin biosynthesis, polar transport, and signaling are well investigated in LR formation. Auxin transport is mediated by efflux carriers PIN-FORMEDs (PINs), ATP-binding cassette Bs (ABCBs), and influx carriers AUX1/LAXs.^10–14^ The auxin transport quadruple mutant *pin1 pin3 pin4 pin7* shows severe defects in LR formation.^15, 16^ Loss of *LAX3* disrupts LR emergence, and the *aux1 lax3* double mutant exhibits an LR initiation defect.^14, 17^ Auxin signaling modules consisting of Aux/IAA transcriptional repressors and auxin response factors drive LR initiation by derepressing ARF7/ARF19 to activate the expression of *LBD16*, *LBD18*, *LBD29*, and *LBD33*.^18–20^

ABA is a well-known stress hormone that plays a central role in the response to environmental stresses in plants and is involved in the regulation of LR development.^21, 22^ ABA is known to suppress LR formation,^23, 24^ which requires the PYR/PYL/RCAR-dependent signaling pathway in *Arabidopsis*,^25^ and inhibition of LR formation occurs at the organ initiation stage.^26^ Moreover, PYL-PP2A signaling modulates LR formation under salt and osmotic stress conditions through increased PIN phosphorylation, thus regulating directional auxin transport.^27^ Notably, the endodermis is required for ABA signaling-mediated LR quiescence when subjected to salt stress.^28^

The coordination between auxin and ABA signaling is necessary for a rapid response to environmental changes but has not yet been well addressed in LR formation. *ABI4*, encoding an ABA-regulated AP2 domain transcription factor, negatively regulates LR number by reducing PIN1 levels in roots.^29^ Ectopic expression of maize *VP1* in the *Arabidopsis abi3* mutant revealed that VP1 mediates a novel interaction between ABA and auxin in roots.^30^ Moreover, the *ABI3* gene is induced in LRP by auxin and ABA and is involved in auxin signaling to mediate LR development in *Arabidopsis*.^31, 32^ Furthermore, *WRKY46*, which is inhibited by ABA but induced by an ABA-independent signal under osmotic/salt stress conditions, directly regulates the expression of *ABI4* and auxin-conjugating genes, contributing to promoting LR formation.^33^ In addition to transcription factors, a RING domain-containing E3 ligase, XBAT32, was identified to modulate LR development by inhibiting ethylene signaling and synergizing the effect of ABA and auxin.^34, 35^ The ABA receptor PYL8 promotes the growth recovery of LRs by interacting with the transcription factors MYB77, MYB44, and MYB73 to mediate the synergistic action of ABA and auxin signaling.^36^

The stress-responsive transcription factor ERF1 acts as a key integrator of environmental signals in *Arabidopsis*.^37–39^ We recently reported that ERF1 promotes local auxin accumulation with altered distribution to inhibit LR emergence by upregulating *PIN1* and *AUX1* and repressing *ARF7* expression.^40^ In this study, we show that the ABI3-ERF1 module mediates ABA-auxin crosstalk to regulate LR emergence. ABA activates *ERF1* expression in LR development, thereby upregulating ERF1-mediated auxin accumulation with altered distribution. Moreover, ABI3 functions as a negative factor to modulate LR formation. ERF1 transcriptionally activates *ABI3*, and reciprocally ABI3 activates *ERF1*. When the protein levels of ABI3 and ERF1 increase, ABI3 interacts with ERF1, and the ABI3-ERF1 complex reduces the ERF1-regulated expression of *PIN1*, *AUX1*, and *ARF7* and ABI3-regulated *ABI5* by decreasing their *cis* element-binding activity, which may serve as a molecular rheostat to prevent auxin and ABA signaling from overamplification.

Therefore, our study provides new insights on the crosstalk between ABA and auxin signaling in LR emergence where the ABI3-ERF1 module plays a crucial role.

## Results

### ERF1 mediates ABA signaling to inhibit LR emergence

To investigate the role of *ERF1* in LR emergence in response to ABA, we examined the expression of *ERF1* in the roots of 7-day-old wild-type (WT) seedlings by qRTLJPCR. The transcript level of *ERF1* was significantly enhanced and peaked at 3 hours after ABA treatment (Figure S1A). Meanwhile, the response of *ERF1* to ABA was examined in the *ERF1pro::GUS* line. GUS staining revealed that *ERF1* expression was significantly induced by ABA in LRP at stages I-VII (Figure S1B), consistent with the qRTLJPCR results. These results suggest that *ERF1* may mediate ABA signaling to regulate LR emergence.

To determine whether ABA affects *ERF1*-mediated LR emergence, we used the RNAi knockdown line *RNAi-1*, the knockout mutant *erf1-2* generated by the CRISPRLJCas9 system, and overexpression lines (OX-2 and OX-4) we previously used^39, 40^ for LR development assays. We found that *RNAi-1* and *erf1-2* mutants exhibited a longer primary root, whereas the overexpression lines showed a shorter primary root than the WT (Figures 1A and 1B), consistent with our previous report.^39, 40^ The primary root elongation of the mutants was less sensitive to ABA treatment than that of the WT and OX lines (Figure 1B). Meanwhile, *RNAi-1* and *erf1-2* mutants displayed more LRs than WT under mock treatment (0 μM ABA), while overexpression lines exhibited much fewer LRs. When subjected to ABA treatment, the *RNAi-1* and *erf1-2* mutants exhibited insignificant changes in LR number and LR density compared with the control, whereas the WT and OX lines showed significant decreases in an ABA dose-dependent manner (Figures 1C and 1D). Moreover, we examined the total lateral root primordium (LRP) number and density of all three *ERF1* genotypes in response to ABA. The *RNAi-1* and *erf1-2* mutants showed no significant changes in LRP number and LRP density under all ABA concentrations compared with the control, whereas the WT and OX lines showed significant decreases in an ABA dose-dependent manner (Figures 1E and 1F). These results indicate that *ERF1* is involved in the inhibition of LR emergence by ABA.

**Figure 1.**
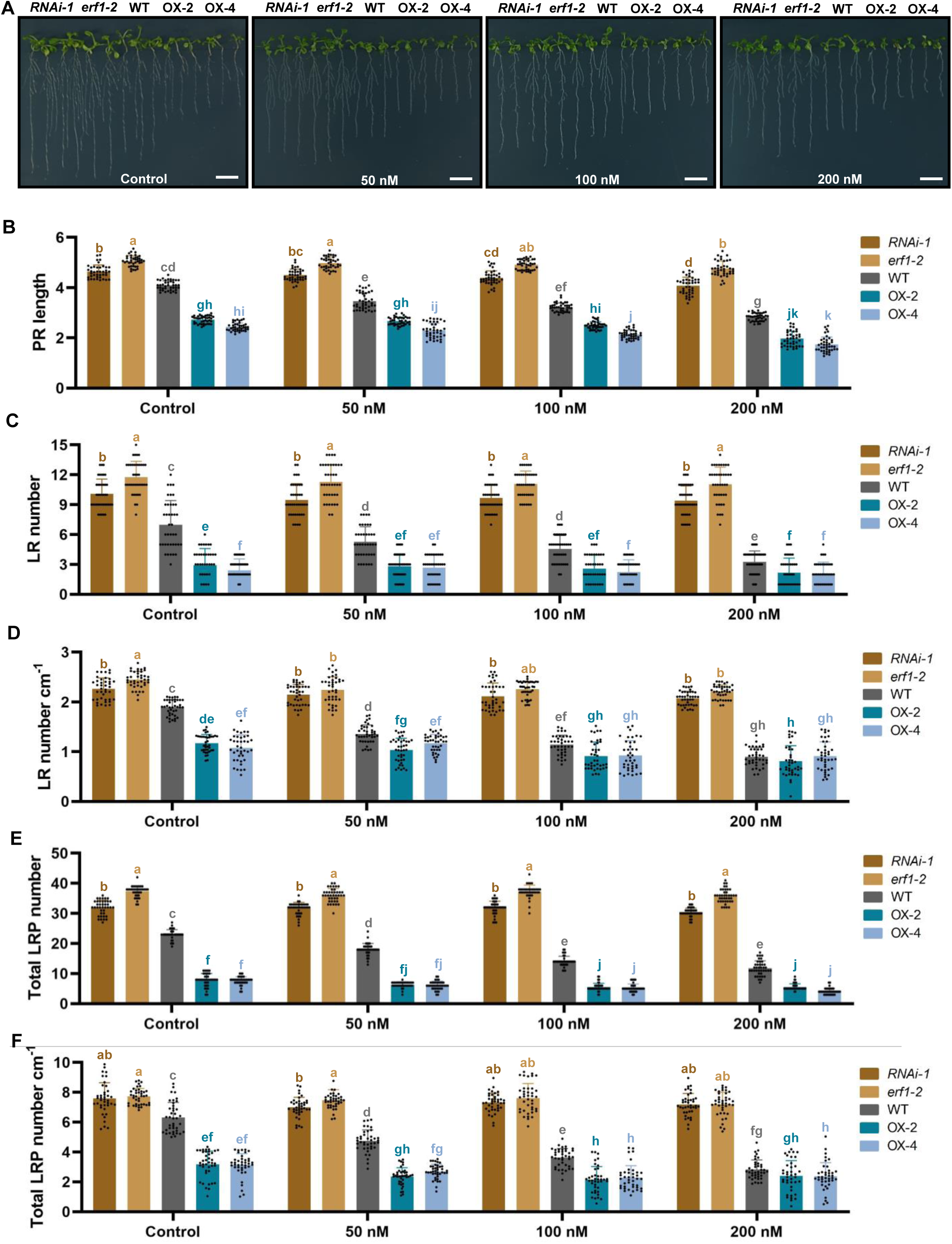
ERF1 mediates LR emergence in response to ABA. (A-D) Sensitivity of LR emergence to ABA. Seeds of different *ERF1* genotypes (knockdown mutant: *RNAi-1*; knockout mutant: *erf1-2*; wild type; overexpression lines: OX-2 and OX-4) were germinated on MS medium for 7 days, and then the seedlings were transferred to MS medium with 0, 50, 100, or 200 nM ABA and vertically grown for 5 days. Photographs were taken (A). Primary root length (B) and LR number (C) were counted, and LR number cm^-1^ was calculated at day 5 (D). Values are the mean ± SD (n=3 replicates, 40 seedlings/replicate). Different letters indicate significant differences by one-way ANOVA (*P* < 0.05). Bar, 1 cm. (E and F) The total LRP number at stages I-VIII. The seeds of different *ERF1* genotypes were germinated and vertically grown on MS medium for 7 days, and then the seedlings were transferred to MS medium with 0, 50, 100, or 200 nM ABA for 2 days. The total LRP number (E) at stages I-VIII was counted, and LRP number cm^-1^ (F) was calculated. Values are the mean ± SD (n=3 replicates, 40 seedlings/replicate). Different letters indicate significant differences by one-way ANOVA (*P* < 0.05). See also Figure S1.

**Figure 2.**
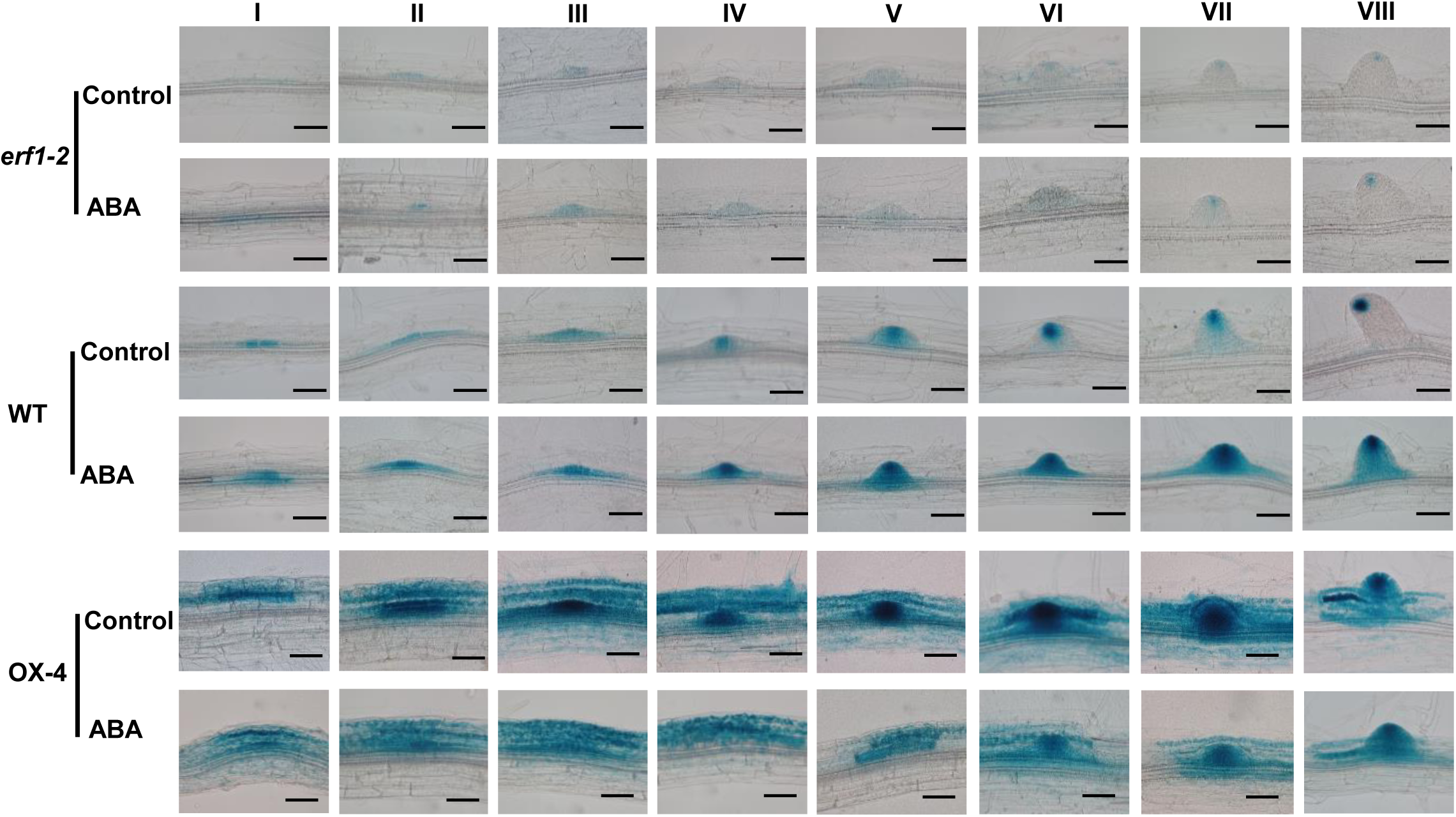
ABA promotes ERF1-mediated auxin accumulation in LRP. Seeds of the *erf1-2*, WT, and OX-4 lines in the *DR5::GUS* background were germinated on MS medium for 5 days, and then the seedlings were transferred to MS medium with or without 200 nM ABA and vertically grown for 2 days before GUS staining for 3 hours. Photographs of stages I-VIII of LR development were taken with a microscope. Values are the mean ± SD (n=3 replicates, 40 seedlings per replicate). Scale bars, 50 μm. See also Figures S2 and S3.

### ABA regulates ERF1-mediated auxin accumulation

To examine whether ABA affects auxin accumulation in an *ERF1*-dependent manner, we histologically observed the expression levels of auxin-responsive *DR5::GUS* in the LRP of *erf1-2*, WT, and OX-4 lines background. Under normal conditions, *erf1-2* exhibited weak GUS signals in the LRP compared with WT, whereas the OX-4 line showed strong GUS signals in the epidermis, cortex, and endodermis cells overlying the LRP (Figure 2), which is consistent with the GUS pattern reported by Zhao et al..^40^ ABA treatment significantly increased GUS signals in the WT line, but the induced GUS signals were abolished in the *erf1-2* line. However, GUS signals were unexpectedly decreased by ABA in OX-4 line. These results suggest that ABA regualtes ERF1-mediated auxin accumulation in the region where LRP initiate. The unexpected decrease in *DR5::GUS* expression in response to ABA suggest that other regulatory mechanisms are involved.

Recently ERF1 was demonstrated to inhibit LR emergence by promoting local auxin accumulation with altered distribution by upregulating *PIN1* and *AUX1* and repressing *ARF7* expression.^40^ To explore whether ABA-attenuated auxin accumulation and inhibition of LR emergence in OX-4 line follows the same mechanism mediated by ERF1, we examined the transcript level of *PIN1*, *AUX1*, and *ARF7* in *erf1-2*, WT, and OX-4 lines in the absence or presence of exogenous ABA. Under no ABA treatment (Control), *PIN1* and *AUX1* were downregulated in *erf1-2* mutant and significantly upregualted in OX-4 line (Figures S2A and S2B), while the transcript level of *ARF7* in OX-4 line was reduced compared with that in WT, but significantly enhanced in *erf1-2* mutant (Figure S2C). Under ABA treatment, the expression levels of *PIN1* and *AUX1* in OX-4 line were lower than those under control treatment, while the transcript abundance of *ARF7* was higher than that in control (Figure S2A-2C). Likewise, the expression patterns of *PIN1pro::GUS* (Figure S2D), *AUX1pro::GUS* (Figure S2E), *ARF7pro::GUS* (Figure S2F) in response to ABA were consistent with qRTLJPCR results. These data suggest that the inhibition of LR emergence by ABA follows the same mechanism by which ERF1 meidates high auxin accumulation with altered distribution by directly regulating *PIN1*, *AUX1*, and *ARF7* as previously reported.^40^ However, the lower expression levels of *PIN1* and *AUX1* in OX-4 under ABA treatment compared with control were unexpected, indicating other regulatory mechanisms might be involved.

### ABI3 transcriptionally activates *ERF1* expression

Having demonstrated that ABA enhances *ERF1*-mediated auxin accumulation, we further investigated whether *ERF1* was regulated by ABA signaling. We first examined the LR phenotype of ABA signaling mutants and found that when grown on MS medium, the *abi3-8* mutant exhibited significantly higher LR number and LR density than the WT, while the overexpression lines of *ABI3* (OX1*^ABI^*^3^ and OX3*^ABI3^*) showed fewer LRs and lower LR density. When subjected to ABA treatments, the *abi3-8* mutant exhibited no significant difference in LR number and LR density compared with the control, whereas the WT and overexpression lines showed decreased LR number and LR density in an ABA dose-dependent manner (Figures 3A-3C). These results indicate that *ABI3* is involved in ABA-mediated inhibition of LR emergence.

**Figure 3.**
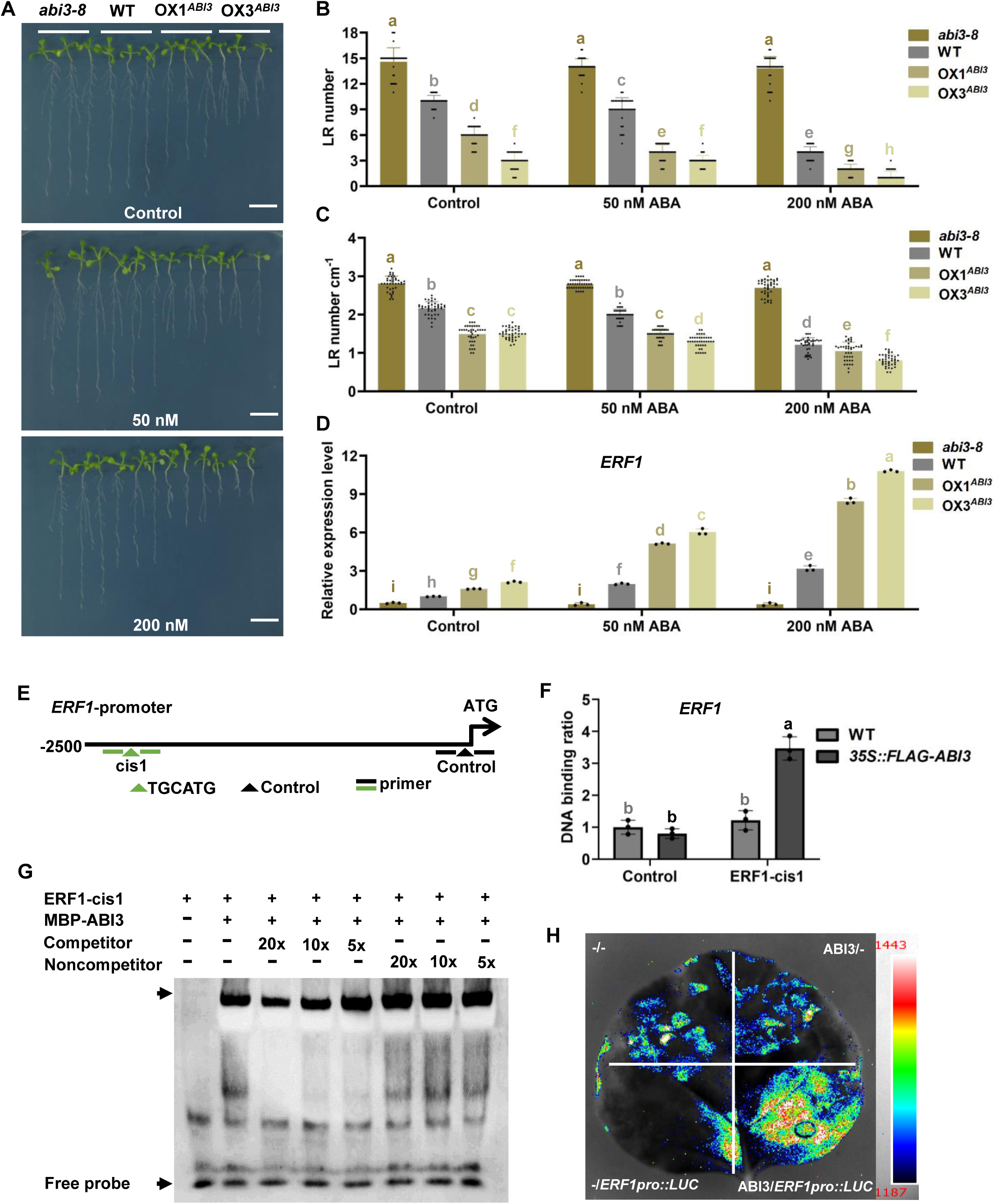
ABI3 negatively regulates LR emergence and activates *ERF1*. (A-C) *ABI3* reduces LR emergence. Seeds of different *ABI3* genotypes (*abi3-8*; WT; *35S::Flag-ABI3* overexpression lines: OX1*^ABI3^* and OX3*^ABI3^*) were germinated on MS medium for 7 days, and then seedlings were transferred to MS medium with 0, 50, or 200 nM ABA for 5 days. Photographs were taken (A). The LR number was counted (B), and the LR number cm^-1^ was calculated at day 5 (C). Values are the mean ± SD (n=3 replicates, 40 seedlings per replicate). Different letters indicate significant differences by one-way ANOVA (*P* < 0.05). Bar, 1 cm. (D) Expression pattern of *ERF1* in the *abi3* mutant and ABI3-OX lines. Seeds of different *ABI3* genotypes were germinated on MS medium for 7 days and transferred to MS medium with 0, 50, or 200 nM ABA for 2 days. Total RNA was isolated from roots for qRTLJPCR analysis. The relative expression level of *ERF1* was normalized to the WT of the corresponding untreated samples. Values are the mean ± SD (n = 3 experiments). (E) Schematic illustration of the location of the RY motif (TGCATG) in the promoter of *ERF1*. The location of the RY motif is indicated with green triangles, while the black triangle indicates a control without the RY motif in the *ERF1* promoter. These lines represent the region amplified in qPCR. (F) ChIP-PCR assay. Ten-day-old wild-type (WT) and *35S::Flag-ABI3* transgenic seedlings grown on MS medium were used for the ChIP-PCR assay. An approximately 200 bp *ERF1* promoter fragment (cis1) containing the RY motif was enriched by anti-FLAG antibodies in qPCR analyses. The region of the *ERF1* promoter that does not contain the RY motif was used as a control. Values are the mean ± SD (n=3 replicates). Different letters indicate significant differences by one-way ANOVA (*P* < 0.05). (G) EMSA. Biotin-labeled *ERF1*-binding cis1 was used as a probe. An unlabeled probe was used as a competitor, and a mutated probe was used as a noncompetitor. “+” represents the presence of the component. “-” represents the absence of the component. As indicated, ABI3-dependent mobility shifts were competed by the competitor probe in a dose-dependent manner (5 ×, 10 ×, 20 ×). Similar results were obtained from three repeat experiments. (H) Transient expression assay in tobacco leaves. The different plasmid combinations (-/- indicates pRI101 and pGreen0800 plasmids, ABI3/- indicates pRI101/*ABI3* and pGreen0800 plasmids, -/*ERF1pro::LUC* indicates pRI101 and pGreen0800/*ERF1pro::LUC*, ABI3/*ERF1pro*::*LUC* indicates pRI101/*ABI3* and pGreen0800/*ERF1pro*::*LUC* plasmids) were transformed into leaf cells of *N. benthamiana* leaves by agroinfiltration. The relative LUC fluorescence intensity was determined by the luciferase assay system (Tannon 5200 M) at 1 day after leaf infiltration using XenolightTM d-luciferin potassium salt. See also Figures S4.

To confirm that ABA signaling is involved in ERF1-modulated LR emergence, we examined the transcript level of *ERF1* in the *abi3-8* mutant and ABI3-OX lines in the absence or presence of exogenous ABA. As shown in Figure 3D, *ERF1* expression was induced in the WT and OX lines by ABA. However, the induced expression of *ERF1* was abolished in the *abi3-8* mutant. These results indicate that ABI3 positively regulates *ERF1* expression.

To address whether ABI3 transcriptionally regulates *ERF1*, we found a potential binding motif (TGCATG) of ABI3 in the 2.5 kb promoter region of *ERF1* (Figure 3E). ChIPLJqPCR assays using 10-day-old WT and *35S::Flag-ABI3* plants revealed a significant enrichment of fragments with the binding motif (Figure 3F). In addition, ABI3 was able to directly bind to the promoter of *ERF1 in vitro,* as shown by electrophoretic mobility shift assay (EMSA) (Figure 3G). Consistent with these results, transient assays in *Nicotiana benthamiana* leaves showed that ABI3 activated *ERF1* expression (Figure 3H). Together, these results demonstrate that ABI3 can directly upregulate *ERF1* expression.

### ERF1 reciprocally activates *ABI3* expression

To search for possible targets of ERF1 in ABA signaling, we checked the expression of ABA signaling-related genes in *erf1-2*, WT, and OX-4 lines by qRTLJPCR analysis. Under normal conditions, *ABI1*, *ABI2*, *ABI3*, *ABI4*, and *ABI5* were significantly upregulated in the OX-4 line and downregulated in the *erf1-2* line, except *ABI1* and *ABI5*, compared with those of WT (Figure S3). In the presence of ABA, the transcript levels of *ABI1*, *ABI2*, *ABI3*, *ABI4*, and *ABI5* were higher in the OX-4 line than in the WT and lower in the *erf1-2* mutant. Additionally, the transcription of *ABI2*, *ABI3* and *ABI5* was not induced by ABA in the *erf1-2* mutant. These results suggest that *ABI3* may be a target gene of ERF1.

To determine whether ERF1 transcriptionally regulates *ABI3*, we searched the potential binding site (GCC-box) of ERF1 in the 2.5 kb promotor and coding region of *ABI3* and found one binding site in the coding region. ChIPLJqPCR, EMSA, and transient expression assays proved the binding and transcriptional activation *in vivo* as shown in Figures 4A-4D. These results demonstrate that ERF1 is able to activate the transcription of *ABI3* by binding to the GCC box in its promoter.

**Figure 4.**
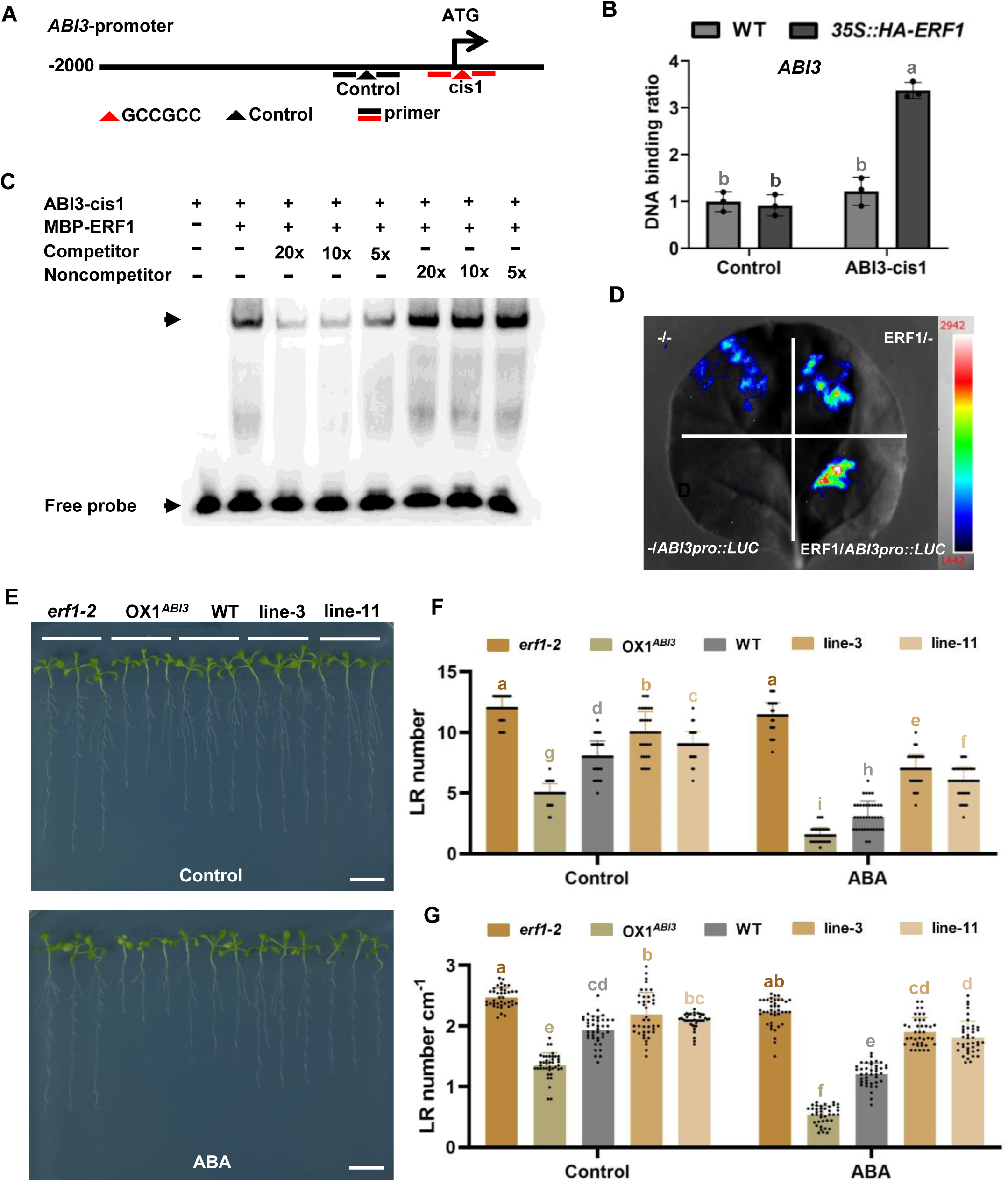
ERF1 activates *ABI3* expression. (A) Schematic illustration of the location of GCC-box (GCCGCC) in the coding region of *ABI3*. The location of the GCC box is indicated with a red triangle, while the black triangle indicates a control without the GCC box motif in the *ABI3* promoter. These lines represent the region amplified in qPCR. (B) ChIP-PCR assay. Ten-day-old wild-type and *35S::HA-ERF1* transgenic seedlings grown on MS medium were used for the ChIP-PCR assay. Approximately 200 bp *ABI3* promoter fragments (cis1) containing the GCC box were enriched by anti-HA antibodies in qPCR analyses. The region of the *ABI3* promoter that does not contain the GCC box was used as a control. Values are the mean ± SD (n=3 replicates). Different letters indicate significant differences by one-way ANOVA (*P* < 0.05). (C) EMSA. Biotin-labeled *ABI3*-binding cis1 was used as a probe. An unlabeled probe was used as a competitor, and a mutated probe was used as a noncompetitor. “+” represents the presence of the component. “-” represents the absence of the component. As indicated, ERF1-dependent mobility shifts were detected and competed with the unlabeled probes in a dose-dependent manner. Similar results were obtained from three repeat experiments. (D) Transient expression assay in tobacco leaves. The different plasmid combinations (-/- indicates pRI101 and pGreen0800 plasmids, ERF1/- indicates pRI101/*ERF1* and pGreen0800 plasmids, -/*ABI3pro::LUC* indicates pRI101 and pGreen0800/*ABI3pro::LUC*, ERF1/*ABI3pro::LUC* indicates pRI101/*ERF1* and pGreen0800/*ABI3pro::LUC* plasmids) were transformed into leaf cells of *N. benthamiana* leaves by agroinfiltration. The relative LUC fluorescence intensity was determined by the luciferase assay system (Tannon 5200 M) at 1 day after leaf infiltration using XenolightTM d-luciferin potassium salt. (E-G) LR emergence of *erf1-2*, WT, OX1*^ABI3^*, line-3 (*erf1-2* in OX1*^ABI3^*, #3), and line-11 (*erf1-2* in OX1*^ABI3^*, #11). Seeds were germinated on MS medium for 7 days, then the seedlings were transferred to MS medium with or without 200 nM ABA and grown vertically for 5 days before photographs were taken (E) and the LR number was counted (F). The LR number cm^-1^ was calculated at day 5 (G). Values are the mean ± SD (n=3 replicates, 40 seedlings/replicate). Different letters indicate significant differences by one-way ANOVA (*P* < 0.05). See also Figures S3 and S4.

We further confirmed the transcriptional regulation of *ABI3* by ERF1 in *ABI3pro::GUS* reporter lines with different *ERF1* backgrounds (*erf1-2*, WT, and OX-4). The expression of *ABI3pro::GUS* was significantly increased in the OX-4 line but reduced in the *erf1-2* mutant compared with the WT without ABA treatment (Figure S4), whereas with ABA treatment, the GUS signals were significantly increased in the WT and OX-4 lines but showed no apparent changes in the *erf1-2* mutant. Taken together, these results indicate that ERF1 positively regulates the transcription of *ABI3*.

To genetically verify the relationship between *ABI3* and *ERF1*, we generated the double mutant OX1*^ABI3^ erf1-2* (line-3 and line-11) by crossing *35S::Flag-ABI3* line with the *erf1-2* mutant. The root phenotype of the double mutants was more similar to that of *erf1-2,* with increased LR number, LR density and primary root length in the absence or presence of ABA treatment compared with that of the OX1*^ABI3^* line (Figures 4E-4G), indicating that *ABI3*-mediated inhibition of LR emergence is dependent on *ERF1* and that *ABI3* is genetically epistatic to *ERF1*.

### ABI3 physically interacts with ERF1 and attenuated the transcription of ERF1/ABI3-regulated genes

The unexpected lower expression levels of *DR5::GUS*, *PIN1* and *AUX1* in OX-4 under ABA treatment (Figure 2 and S2) promoted us to explore other regulatory mechanisms. We speculated that there might be a faster response by proteinLJprotein interaction between ABA and auxin signaling components and searched for ERF1-interacting partners by yeast two-hybrid assays. Interestingly, we found that ABI3 and ERF1 physically interacted in yeast cells (Figure 5A). To confirm this interaction, we performed a GST pull-down assay and found that GST-ERF1, but not GST, interacted with HIS-ABI3 *in vitro* (Figure 5B). Moreover, we further confirmed this interaction with coimmunoprecipitation (Co-IP) and bimolecular fluorescence complementation (BiFC) assays. In the Co-IP assay, Flag-ABI3 was immunoprecipitated by HA-ERF1 but not by Flag (Figure 5C). In the BiFC assay, YFP fluorescence was observed only in the nuclei when ERF-nYFP was coinfiltrated with ABI3-cYFP in tobacco leaves, and no fluorescence was observed in the negative controls (Figure 5D). Together, these results clearly demonstrate that ERF1 can interact with ABI3 *in vitro* and *in vivo*.

**Figure 5.**
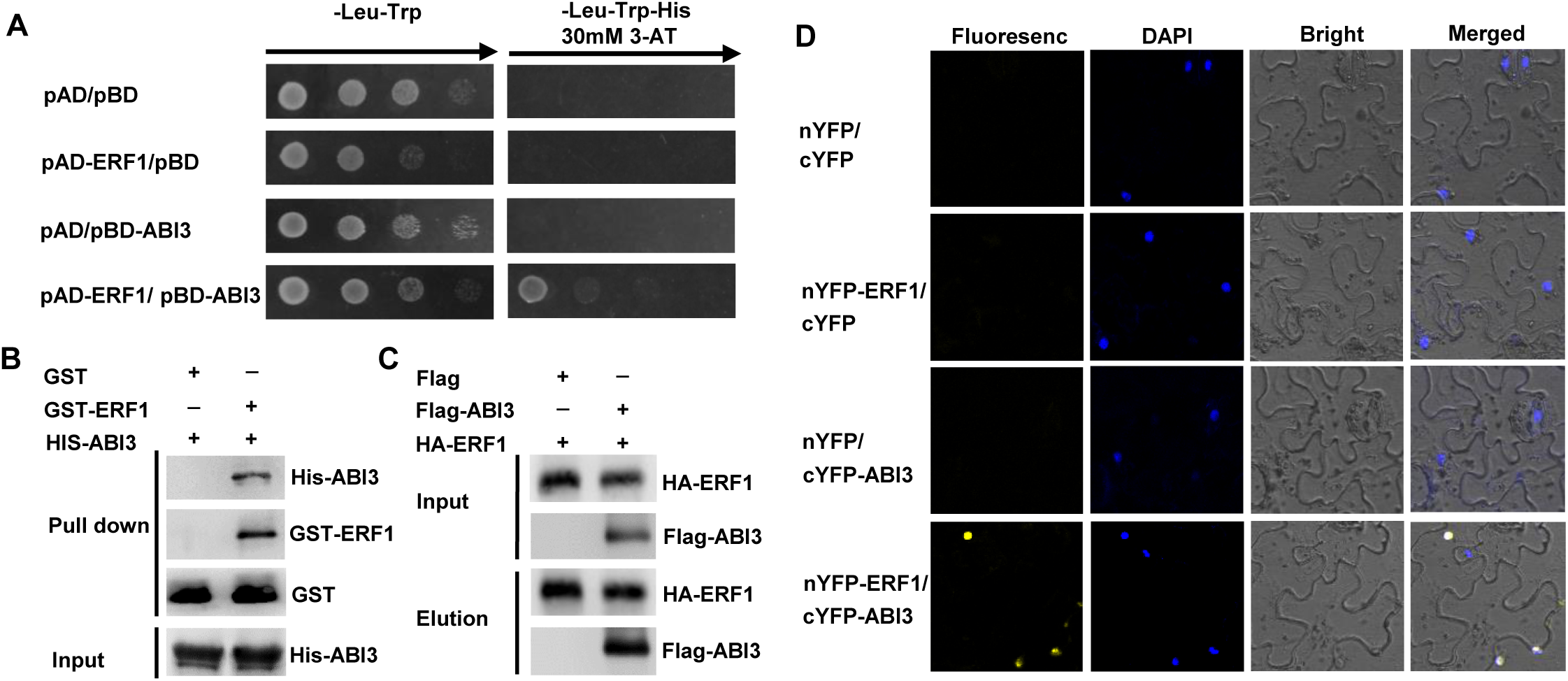
ERF1 interacts with ABI3. (A) Y2H assay. The interaction was indicated by the ability of yeast cells to grow on selective SD-Leu-Trp-His medium with 30 mM 3-AT. pAD/pBD, pAD-ERF1/pBD, and pAD/pBD-ABI3 were used as negative controls. (B) GST pull-down assay. GST and GST-ERF1 were incubated with amylose resin bound to the HIS-ABI3 protein. The pulled-down protein complex was detected by western blot analyses using an anti-His antibody. (C) Co-IP assay. *N. benthamiana* leaves were transfected with plasmids expressing HA-ERF1 and Flag-ABI3/Flag-as indicated by agroinfiltration and incubated for 48 hours. Cell lysates were subjected to Co-IP with anti-HA or anti-Flag antibodies, followed by western blotting with related antibodies. (D) BiFC assay. Different plasmid combinations were expressed in leaf cells of *N. benthamiana* leaves. Yellow fluorescence protein (YFP) was observed in epidermal cells expressing both nYFP-ERF1 (the N-terminal part of YFP fused with ERF1) and cYFP-ABI3 (the C-terminal part of YFP fused with ABI3). nYFP/cYFP, nYFP-ERF1/cYFP, and nYFP/cYFP-ABI3 were used as negative controls. nYFP (pAS-54) indicates the NE vector, and cYFP (pAS-58) indicates the CE vector. DAPI staining marks the nucleus. See also Figures S5.

Given that ERF1 physically interacts with ABI3, we then asked what the biological consequence of the interaction would be. We recently reported that ERF1 can activate the transcription of *PIN1* and *AUX1* and repress *ARF7*.^40^ Therefore we performed EMSA to determine whether the binding activities were altered by the interaction. The results in Figures 6A-6C demonstrate that ABI3 attenuated the DNA binding activity of ERF1 to the *cis* element in the promoter of *PIN1*, *AUX1* and *ARF7*.

**Figure 6.**
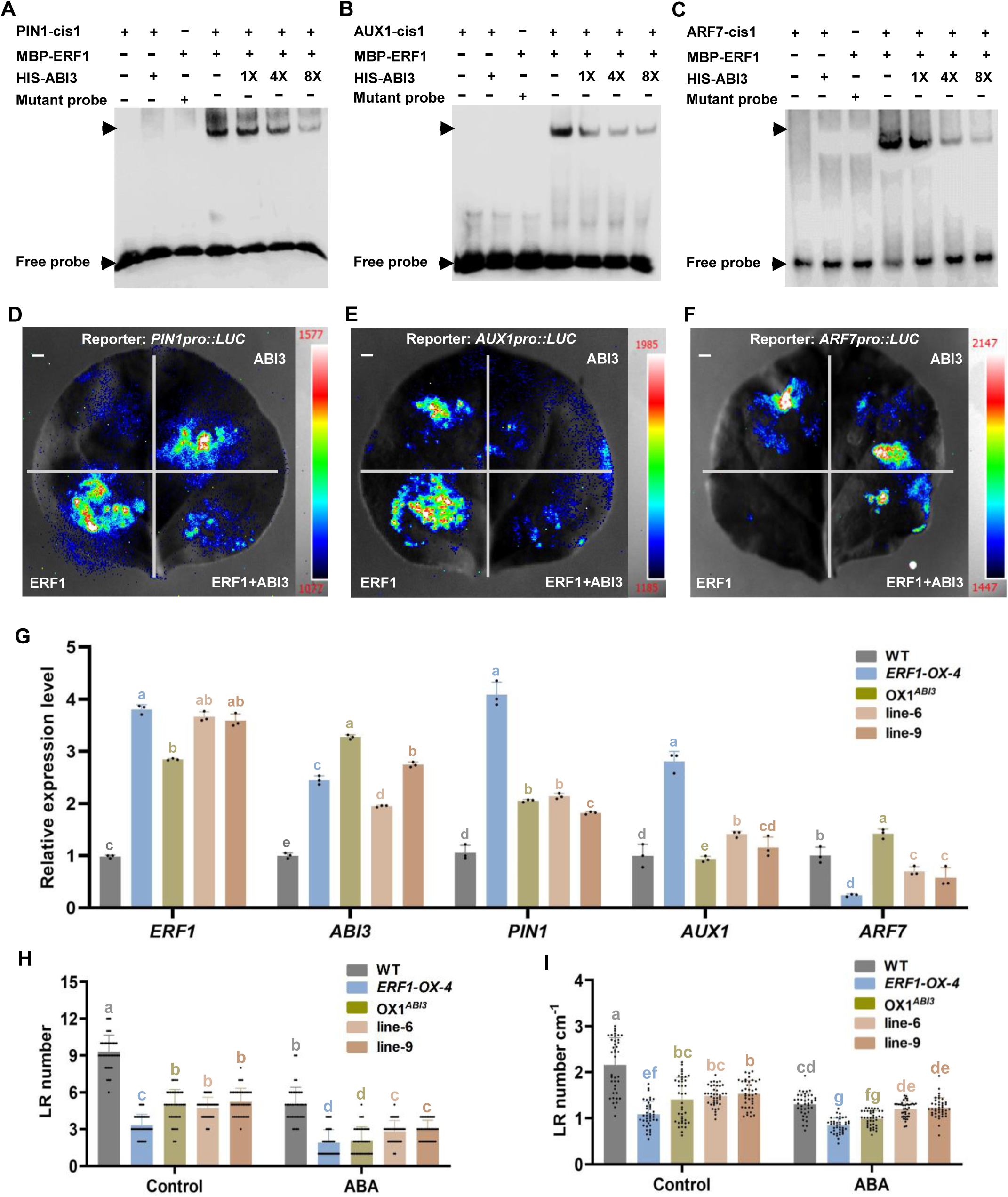
ABI3-ERF1 interaction reduces the binding activity of ERF1 and attenuates the expression of ERF1-regulated genes. (A-C) EMSA. Biotin-labeled *PIN1*-cis1, *AUX1*-cis1, and *ARF7*-cis1 were used as probes. Unlabeled probes were used as competitors, and mutated probes were used as noncompetitors. “+” represents the presence of the component. “-” represents the absence of the component. As indicated, ERF1-dependent mobility shifts were detected and competed with the HIS-ABI3 protein in a dose-dependent manner. Similar results were obtained from three repeat experiments. (D-F) Transient expression assay in tobacco leaves. pRI101-ERF1/ABI3 both act as effectors. pGreenII0800-*PIN1*/*AUX1*/*ARF7* function as reporters. “-’’ indicates pRI101 empty plasmid. The different plasmid combinations were transformed into leaf cells of *N. benthamiana* leaves by agroinfiltration. The relative LUC fluorescence intensity was determined by the luciferase assay system (Tannon 5200 M) at 1 day after leaf infiltration using XenolightTM d-luciferin potassium salt. (G) The transcript levels of ERF1-regulated genes. Seeds of the WT, *ERF1-OX*, line-6 (*ERF1-OX* in *ABI3-OX*, #6), and line-9 (*ERF1-OX* in *ABI3-OX*, #9) lines were germinated on MS medium for 7 days, and then RNA was isolated from roots. The relative expression levels of *ERF1* and *ABI3*, *PIN1*, *AUX1*, and *ARF7* were analyzed by qRTLJPCR and normalized to WT. Values are the mean ± SD (n=3 replicates). Different letters indicate significant differences by one-way ANOVA (*P* < 0.05). (H and I) Overexpression of *ABI3* in the *ERF1-OX* line alleviates the inhibition of LR emergence. Seeds of WT, *ERF1-OX-4*, OX1*^ABI3^*, line-6 (*ERF1-OX-4* in OX1*^ABI3^*, #6), and line-9 (*ERF1-OX-4* in OX1*^ABI3^*, #9) lines were germinated on MS medium for 7 days, and then seedlings were transferred to MS medium with 0 or 200 nM ABA for 5 days. The LR number was counted (H), and the LR number cm^-1^ was calculated at day 5 (I). Values are the mean ± SD (n=3 replicates, 40 seedlings/replicate). Different letters indicate significant differences by one-way ANOVA (*P* < 0.05). Bar, 1 cm. See also Figures S5.

In addition, we performed transient expression assays in *N. benthamiana* leaves to examine the effect of the ERF1-ABI3 interaction on the expression of ERF1-regulated genes (*PIN1*, *AUX1*, *ARF7*). When ERF1 and ABI3 proteins were coexpressed, *PIN1* and *AUX1* promoter-driven LUC expression was significantly reduced compared to that in the presence of ERF1 alone, while *ARF7* promoter-driven LUC expression was increased (Figures 6D-6F).

To further confirm the biological consequence, we generated transgenic lines (line-6 and line-9) expressing both *35S::ERF1* (*ERF1-OX-4*) and *35S::ABI3* (OX1*^ABI3^*) and measured the expression levels of *PIN1*, *AUX1*, and *ARF7* in these lines (Figure 6G). In the *ERF1-OX-4* line, the transcript abundance of *PIN1* and *AUX1* was enhanced compared with that of WT, and *ARF7* expression was downregulated, whereas in the OX1*^ABI3^ ERF1-OX-4* lines, the expression levels of *PIN1* and *AUX1* were lower than those in *ERF1-OX-4* plants, and the transcript abundance of *ARF7* was higher than that in *ERF1-OX-4* plants (Figure 6G). Furthermore, we analyzed the LR number and density in the WT, *ERF1-OX-4*, OX1*^ABI3^*, and OX1*^ABI3^ ERF1-OX-4* lines and found that the OX1*^ABI3^ ERF1-OX-4* lines (line-6 and line-9) displayed a higher LR number and density than *ERF1-OX-4* but a lower LR number and density than WT in the absence or presence of ABA (Figures 6H and 6I).

Furthermore, we examined whether the ABI3-ERF1 interaction affects the transcription of ABI3-regulated genes (e.g. *ABI5*) by EMSA and transient expression assays in *N. benthamiana*. As shown in Figures S5A and S5B, when ERF1 and ABI3 proteins were coexpressed, the DNA binding activity of ABI3 to *ABI5* promoter and *ABI5* promoter-driven LUC expression were decreased compared with that in the presence of ABI3 protein alone. Likewise, the transcription abundance of *ABI5* was analyzed in *35S::ERF1* (*ERF1-OX-4*), *35S::ABI3* (OX1*^ABI3^*) and the transgenic line-6 and line-9. The expression level of *ABI5* in line-6 and line-9 were lower than those in OX1*^ABI3^* plants. These data illustrated that the ERF1-ABI3 interaction also alters the expression of genes assoiated with ABA signaling.

Taken together, these results demonstrate that the ERF1-ABI3 interaction attenuates ERF1 and ABI3 binding activity to their *cis* element, thereby decreasing the transcription of their target genes and alleviating the inhibitory effects of LR emergence by ERF1 and explain the unexpected lower expression levels of *PIN1* and *AUX1* in OX-4 under ABA treatment compared with control (Figure S2).

## Discussion

### The ABI3-ERF1 module mediates the crosstalk between ABA and auxin to regulate LR emergence

LR development is key for plant root architecture to satisfy survival strategies in response to environmental cues. In our previous report, we demonstrated that stress-responsive ERF1 mediates high-level auxin accumulation and inhibition of LR emergence by upregulating the expression of *PIN1* and *AUX1* to enhance auxin transport and downregulating the expression of *ARF7*, which leads to the altered expression of cell wall remodeling genes, by which plants adjust their root system architecture to adapt to changing environments.^40^ In this study, we identified the ABI3-ERF1 module that integrates ABA signaling to enhance ERF1-mediated auxin accumulation and inhibit LR emergence. A working model is proposed (Figure 7).

**Figure 7.**
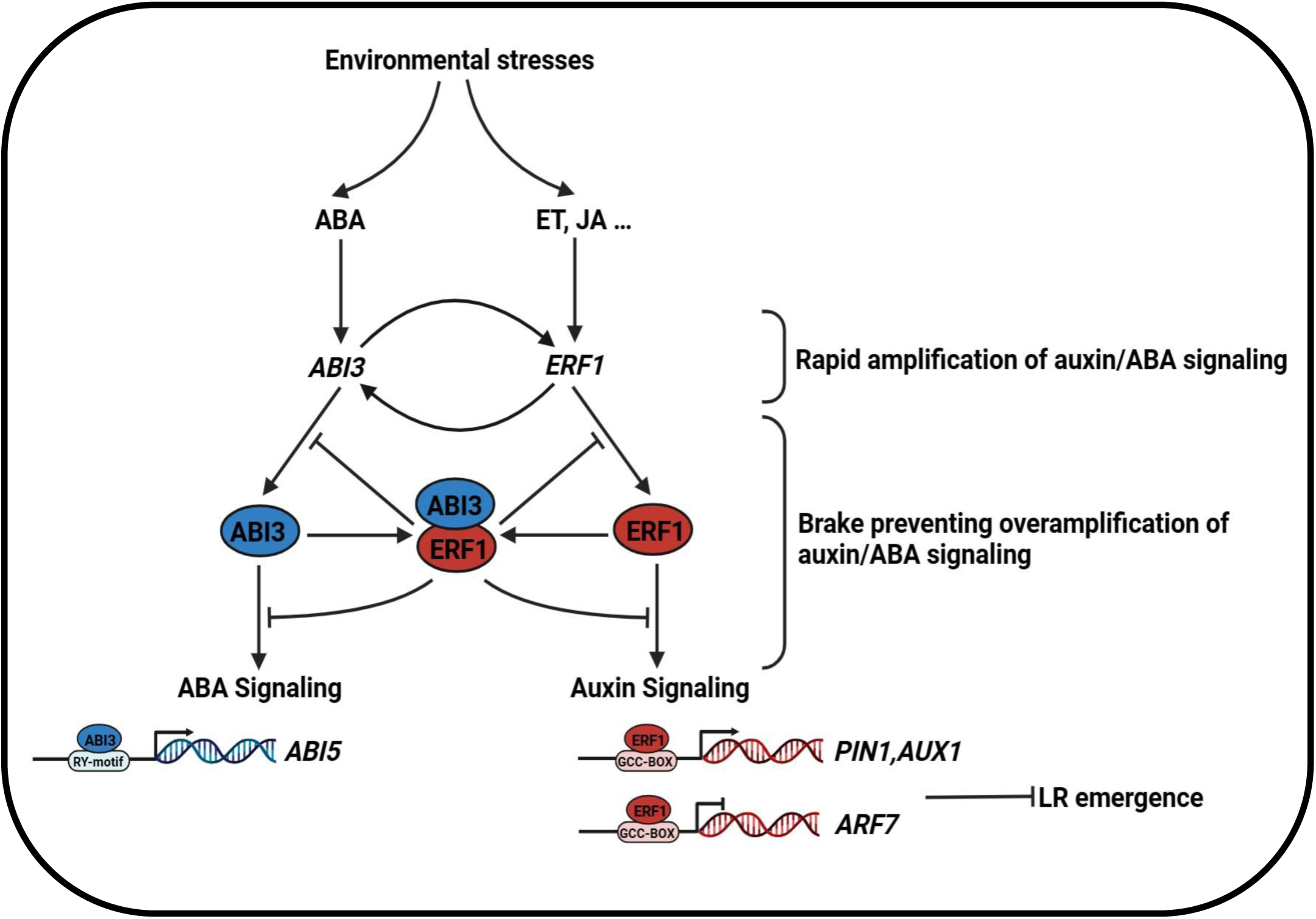
A working model for the ABI3-ERF1 module in mediating ABA-auxin crosstalk in LR emergence. At the transcriptional level, ABI3 transcriptionally activates *ERF1* in response to ABA. Reciprocally, ERF1 transcriptionally activates *ABI3*. This coregulation allows rapid amplification of auxin and ABA signaling. When ABI3 and ERF1 proteins reach certain levels, they physically interacts with each other, decreases their binding ability, thereby attenuates ERF1-mediated transcriptional regulation of genes such as *PIN1*, *AUX1* and *ARF7*, which modulate LR emergence, as demonstrated in our previous report,^40^ and ABI3-mediated transcriptional regulation of *ABI5*. Therefore, ABI3-ERF1 complex may serve as a molecular rheostat to avoid overamplification of auxin and ABA signaling in LR development and fine-tune the crosstalk between ABA and auxin signaling. See also Figures S4, and S5.

Under stress conditions that trigger ABA signaling, ABI3 transcriptionally activates *ERF1*, and reciprocally, *ABI3* is activated by ERF1, rapidly amplifying signals to regulate root development. Moreover, ABI3 can physically interact with ERF1 when their proteins reach certain levels, and this interaction reduces the binding activity of ERF1 and ABI3 and attenuates the transcription of ERF1-regulated genes (*PIN1*, *AUX1*, *ARF7*) and ABI3-regulated gene (*ABI5*), acting as a brake to prevent overamplification of auxin and ABA signaling. This ERF1-centered framework mediates ABA-auxin crosstalk and plays an important role in LR emergence.

### Reciprocal transcriptional upregulation between ERF1 and ABI3 facilitates rapid signal amplification

ERF1 is known to integrate JA, ethylene and auxin signaling to modulate plant growth.^37–39, 41^ To date, it has been reported that the transcript level of *ERF1* in 2-week-old wild-type plants is enhanced by JA and ethylene, while the induction is suppressed by ABA.^38^ However, we showed that ERF1 was induced by exposure to ABA in the roots of the 7-day-old wild type and in the LRP of the 7-day-old *ERF1pro::GUS* transgenic line (Figure S1). This discrepancy in *ERF1* expression in response to ABA may be due to different spatiotemporal expression patterns in plants of different ages.

ABA is known to suppress LR formation and growth.^23, 24, 42^ ABI4, a key regulator in the ABA signaling pathway, is induced in roots by ABA and cytokinin and integrates ABA and cytokinin to inhibit LR formation by reducing the level of PIN1 in *Arabidopsis*.^29^ In addition, ABI3 is also involved in modulating root development.^30, 43, 44^ In our study, we showed that *ABI3* negatively regulates LR emergence (Figures 3A-3C). The induced expression of *ERF1* by ABA was abolished in the *abi3-8* mutant compared with that of WT (Figure 3D), indicating that *ERF1-* mediated LR emergence in response to ABA was dependent on *ABI3*. In particular, ABA signaling components, including *ABI1*, *ABI2*, *ABI3*, *ABI4*, and *ABI5,* were significantly upregulated in the OX-4 line and downregulated in the *erf1-2* mutant compared with the WT under ABA treatment (Figure S3), suggesting that ERF1 can affect ABA signaling to modulate LR formation. Our biochemical and genetic evidence demonstrated that ABI3 activates the transcription of *ERF1* (Figures 3E-3H), and reciprocally ERF1 can directly upregulate *ABI3* expression (Figures 4A-4D and Figure S4). The reciprocal activation between ABI3 and ERF1 forms a positive regulatory loop to amplify auxin and ABA signaling, which may be required to coordinate the crosstalk between ABA-auxin signaling to balance the stress response with plant growth, improving plant survival.

To adapt to changes in the local environment, plants respond with fine regulation to satisfy the needs of growth. For example, the B-BOX (BBX)-containing proteins BBX24/BBX25 regulate *BBX22* transcription by suppressing *HY5* transcriptional activity, thus modulating light-mediated plant development.^45^ Meanwhile, BBX22 physically interacts with HY5, and HY5 also activates the expression of *BBX22*, coregulating hypocotyl growth.^45–47^ Furthermore, ARF17, ARF6 and ARF8 regulate each other’s expression and maintain transcript homeostasis to regulate adventitious rooting in *Arabidopsis*.^48^ Recently, the WRKY transcription factors WRKY33 and WRKY12 were shown to synergistically activate *RAP2.2* transcription, whereas RAP2.2 feedback activates *WRKY33*, thus establishing hypoxia signaling and enhancing hypoxia tolerance.^49^ These complex reciprocal regulatory networks among transcription factors may allow immobile plants to rapidly respond and modulate their adaptive response. This ERF1-centered regulatory mechanism rapidly adjusts at the transcription level to establish ABA-auxin links, ultimately affecting LR emergence.

### The ABI3-ERF1 interaction may serve as a molecular rheostat to avoid overamplification of auxin and ABA signaling

ABI3 acts as a transcriptional coactivator by interacting with ABA-response element (ABRE)-binding factors.^50, 51^ ABI3 mediates the interaction between proteins, especially bZIP factors, by the acidic A1 domain and two other basic domains, B1 and B2.^52, 53^ In this study, we demonstrated that ERF1 physically interacts with ABI3 *in vitro* and *in vivo* (Figure 6), providing another layer of regulation for the crosstalk between ABA and auxin signaling.

ABA treatment increased auxin accumulation in both WT and OX-4 lines during LRP development but not in the *erf1-2* mutant (Figure 2), suggesting that ERF1 was able to play an important role in the coordinated action of ABA and auxin in regulating LR emergence. We reported that ERF1 inhibits LR emergence by enhancing local auxin accumulation with altered distribution by activating *PIN1* and *AUX1* and repressing *ARF7* expression.^40^ Our results showed that the interaction between ABI3 and ERF1 weakened the ERF1-mediated transcriptional regulation of *PIN1*, *AUX1*, and *ARF7* (Figure 6). When *ABI3* was overexpressed in ERF1-OX plants, it displayed a higher LR number and density compared to that of ERF1-OX plants (Figure 6H), which is consistent with the prediction that the ERF1-ABI3 interaction would alleviate the inhibition of ERF1-mediated LR emergence. ABI5 not only plays a key role in ABA core signaling, but also functions as an integrator of ABA and other phytohormone signaling.^54, 55^ *ABI5* expression was induced in the LR tips after ABA treatment,^56^ indicating that ABI5 may act as a regulator mediating environmental stress to regulate LR development. Notably, ERF1-ABI3 interaction also reduced the binding activity of ABI3 to *ABI5* promoter (Figure S5), causing decreased ABA signaling. These findings illustrate that the ABI3-ERF1 interaction may act as a molecular rheostat to avoid auxin and ABA signaling overamplification in response to hormones and environmental cues, which may be required in the balance between stress response and growth.

Enhancing or attenuating transcription regulation by protein interactions is well known. For example, ABI3 and FUSCA3 (FUS3), which contain homologous DNA-binding B3 domains, exhibit common maturation-related phenotypes.^50, 57^ A previous study also illustrated that FUS3 interacts with LEAFY COTYLEDON2 (LEC2), synergistically activating *YUCCA4* transcription during LR formation.^58^ Furthermore, PYL8 enhanced the transcriptional activity of auxin-responsive genes to promote LR growth by interacting with MYB77.^36^ Meanwhile, the physical interaction between PIF4 and PYL8/9 enhanced PIF4 binding to the *ABI5* promoter but inhibited PIF4-mediated transactivation of *ABI5* in *Arabidopsis*.^59^

Auxin-ABA interactions are crucial to LR development, but the underlying mechanisms are not well understood. Recently, it was shown that the *abi3* mutant is resistant to auxin in promoting LR formation.^31^ Additionally, auxin can rescue the inhibitory effects of ABA on LR elongation but not on LR initiation.^60^ Our results showed that the ERF1-ABI3 interaction enables plants to rapidly respond to environmental stresses and acts as a molecular rheostat to prevent signaling overamplification, which may be required for ABA-auxin crosstalk.

### ABI3-ERF1 module-mediated LR emergence serves plants to adapt to changing environments

To survive unpredictable environmental changes, plants have evolved various sophisticated means to detect environmental cues and respond appropriately.^61^ Root plasticity, as a key root trait for adaptation to different environments, determines shoot growth and plant production. It is well known that the hormone ABA mediates the response to various environmental stresses, coordinating with plant growth and development. In this study, we have shown that ERF1 mediates the crosstalk between ABA and auxin signaling via the ERF1-ABI3 module in regulating LR emergence. These regulatory mechanisms may be required to balance plant growth with the stress response for better survival in the changing environment. Therefore, our findings provide new insights into the mechanism of LR emergence mediated by ABA and auxin coordination.

### Limitations of the study

Because ABA promotes ERF1-mediated enhanced auxin accumulation and altered distribution in epidermis, cortex, and endodermis cells overlying the LRP, it will be interesting to explore whether ABA-mediated endodermal response is invovled in ERF-regulated LR emergence. The role of ERF1 in LR emergence is also coordinated with other regulators, which needs to be investigated.

## Acknowledgments

This study was supported by grants from the National Natural Science Foundation of China (31900230 to P.X.), China Postdoctoral Science Foundation (2020T130634 and 2019M652200 to P.X.) and Youth Innovation Foundation of University of Science and Technology of China (WK2070000186 to P.X.). We thank Dr. Xiangyang Hu, Shanghai University for providing the *35S::ABI3* seeds.

## Author contributions

J.Z., P.Z., and C.X. designed the experiments. P.Z., J.Z., Y.C., L.S., J.M. performed the experiments and data analyses. J.Z. wrote the manuscript. P.Z., S.T., and C.X. revised the manuscript. P.Z. and C.X. supervised the project.

## STAR Methods

### Resource availability

#### Lead contact

Further information and requests for reagents and source data should be directed to and will be fulfilled by the lead contact, Chengbin Xiang.

#### Materials availability

The materials generated in this study are available from the corresponding author.

#### Data and code availability

All data reported in this paper will be shared by the lead contact upon request.

This paper does not report original code.

Any additional information required to reanalyze the data reported in this paper is available from the lead contact upon request.

### Experimental model and subject details

#### Plant materials and growth conditions

*Arabidopsis thaliana* ecotype Columbia-0 (WT) was used in our study. *ERF1pro::GUS*, *35S::ERF1-GFP*, *35S::ERF1* (OX-2, OX-4), *ERF1* knockdown mutant (*RNAi-1*), *ERF1* knockout mutant (*erf1-2*), and *DR5::GUS* were used in this study as previously described.^39^ *ABI3pro::GUS* transgenic lines were constructed by cloning the promoter of *ABI3* into pCAMBIA1391Z with primers (Table S1). The genetic materials were obtained by genetic crossing, where *DR5::GUS* and *ABI3pro::GUS* lines were used as female parents, *erf1-2* and *35S::ERF1* lines were used as male parents. The *ERF1-OX ABI3-OX* lines (line-6 and line-9) were generated by crossing the *35S::ERF1* line (female parent) with the *35S::ABI3* line (male parent).

Surface-sterilized *Arabidopsis* seeds were treated with 10% bleach for 15 min and washed four times with at least sterile water. Seeds were kept at 4°C for 2-4 days in darkness before germination on agar plates containing solid Murashige and Skoog (MS) medium with 1% (w/v) sucrose under long-day conditions (16-h light/8-h dark) at 22-24°C.

#### Method details

##### Analysis of LR number and LR density

The analysis of LR number and density was performed as previously described.^62^ Seven-day-old seedlings grown on MS medium were transferred to MS medium without or with different concentrations of abscisic acid (ABA) and grown vertically for 5 days. The PR length of seedlings was measured using ImageJ, and the number of LRs was counted at the indicated time points.

##### Histochemical GUS staining

Histochemical staining for GUS activity in transgenic plants was conducted as previously described.^63^ Seedlings were incubated in GUS staining solution (1 mg/mL X-glucuronide in 0.1 M potassium phosphate, pH 7.2, 0.5 mM ferrocyanide, 0.5 mM ferricyanide, and 0.1% Triton X-100) at 37°C in the dark for the indicated time. After incubation, seedlings were cleared with a series of ethanol solutions (100%, 50%, 30%). Images were captured using an OLYMPUS IX81 microscope and HiROX (Japan) MX5040RZ.

##### RNA isolation and quantitative real-time PCR analysis

Total RNA was extracted by RNA easy^TM^ Isolation Reagent (Vazyme, #r701-02) from roots. cDNA was prepared by HIScriptRLJRT SuperMix for qPCR (Vazyme, #r323-01) and used for quantitative RTLJPCR. Quantitative RTLJPCR assays were performed with a ChamQ SYBR qPCR Master Mix kit on a real-time PCR system (Vazyme, #q311-02) according to the manufacturer’s instructions. The expression levels of target genes were normalized to *Arabidopsis UBQ5*. All qRTLJPCR experiments were performed with at least three biological replicates. The primers used in this study are listed in Table S1.

##### ChIP-PCR assay

ChIP-PCR assays were conducted essentially as previously described.^64^ Ten-day-old *35S::HA-ERF1* seedlings, *35S::Flag-ABI3*, and WT plants were harvested for ChIP experiments. The enrichment of DNA fragments was used for qPCR using the following primers (Table S1). β*-Tubulin8* was used as a negative control. The results represent the average of at least three independent repeats.

##### Protein expression and purification

The coding sequence of *ERF1* was constructed into the pMAL-C2X and pGEX-6P-1 vectors and transformed into the *Escherichia coli* Rosseta3 strain to generate MBP-ERF1 and GST-ERF1 fusion proteins for EMSA and pull-down assays, respectively. The coding sequence of *ABI3* was constructed into the pET-28a vector and transformed into the *Escherichia coli* Rosseta3 strain, in which the HIS-ABI3 fusion protein was expressed for EMSA and pull-down assays. Fusion proteins were purified as previously described.^65^

##### Electrophoretic mobility shift assay

EMSA was performed as previously described ^66^ using a LightShift™ EMSA Optimization and Control Kit (Thermo Fisher Scientific). The 22 bp free probes containing the binding motif, competitor and noncompetitor were synthesized by a commercial company (Sangon Biotech, Shanghai, China). The binding assay samples were separated in a 6% native polyacrylamide gel in 0.5× TBE buffer. The results were detected using a CCD camera system (Image Quant LAS 4000).

##### Transient transactivation assay

A transient transactivation assay in *Nicotiana benthamiana* leaves was conducted as described previously.^67^ The coding sequences of *ERF1* and *ABI3* were amplified and cloned into the pRI101 vector. Approximately 2500 bp promoters of *ERF1* and *ABI3* were cloned into the pGreenII 0800-LUC vector. All constructs were electroporated into the *Agrobacterium* GV3101 strain by using MicroPulser Electroporation Systems (Bio-Rad, CA, USA) and then coinjected into *Nicotiana benthamiana* leaves with the infiltration solution (10 mM MES, 10 mM MgCl_2_, 150 mM acetosyringone, pH 5.6). Two days after injection, tobacco leaves were sprayed with Luc substrates (Xeno light^TM^ D-luciferin potassium salt, 1 mM), and fluorescence images were taken with a Tanon 5200 automatic chemiluminescence imaging system.

##### GST pull-down assay

GST pull-down assay was carried out as described previously.^68^ GST and GST-ERF1 fusion proteins were incubated with GSH-agarose beads at 4°C for 4 h. The beads were washed with washing buffer (1 x PBS) 3 times. Then, the beads were incubated with HIS-ABI3 at 4°C overnight and washed 3 times. Pulled-down mixtures were used to detect HIS-ABI3 with anti-HIS antibody (ABMART, #M20003) and GST-ERF1 with anti-GST antibody (PROTEINTECH, #HRP-66001) by western blotting.

##### BiFC analysis

pAS-054-ERF1 (the N-terminus of YFP fused with ERF1) and pAS-058-ABI3 (the C-terminus of YFP fused with ABI3) were constructed and transferred to *Agrobacterium* strain C58C1. The different plasmid combinations were transformed into leaf cells of *N. benthamiana* leaves by agroinfiltration. YFP and DAPI signals were observed under excitation wavelengths of 510 and 405 nm, respectively, after leaf infiltration for 2-3 days by a Zeiss 880 confocal laser scanning microscope.

##### Coimmunoprecipitation assay

Protein immunoprecipitation using the *Nicotiana benthamiana* transient expression system was conducted as described previously.^69^ pCAM1300-3×HA-ERF1 and pCAM1300-3×Flag-ABI3 were constructed. Flag-tagged proteins were coexpressed with HA-ERF1 in tobacco leaves by agroinfiltration for 2 days. Samples were ground with lysis buffer (150 mM NaCl, 1 mM EDTA, 50 mM Tris-HCl pH 7.5, 0.2% Triton-X-100, 1% NP40, cocktail, 50 μM MG132). The eluates were subjected to western blotting with anti-Flag and anti-HA antibodies.

##### Quantification and statistical analysis

Statistical analyses were performed by SPSS v16.0 software based on one-way ANOVA. Error bars for all data represent the mean ±SD of at least three replicates, as indicated in the figure legend, and significant differences were determined at P≤0.05.

## Supplemental information

Figure S1. *ERF1* is responsive to ABA during LR development. Related to Figure 1.

Figure S2. The expression levels of the ERF1-targeted genes in response to ABA. Related to Figure 2.

Figure S3. The expression levels of the genes involved in ABA signaling were altered by ERF1. Related to Figure 4.

Figure S4. The expression of *ABI3pro::GUS* was enhanced by ERF1. Related to Figure 4.

Figure S5. ABI3-ERF1 interaction reduces the binding activity of ABI3 to *ABI5*. Related to Figure 6.

Table S1. Primers used in this study. Related to Figures 3, 4, 5, 6, S1, S2, S3, S4, and S5.

