## Supplemental info for "The ABI3-ERF1 module mediates ABA-auxin crosstalk to regulate lateral root emergence"

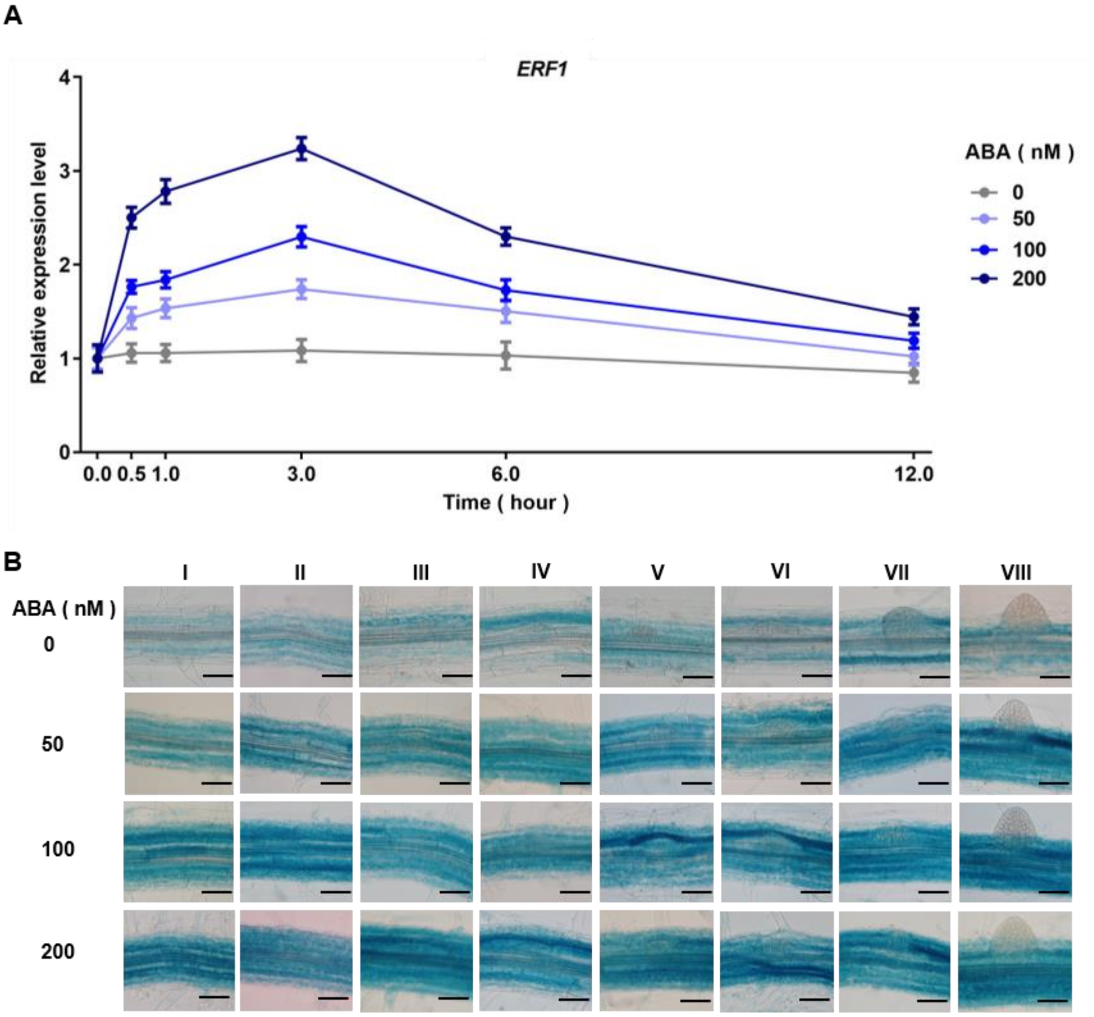


**Figure S1. *ERF1* is responsive to ABA during LR development. Related to Figure 1.**

(A) *ERF1* is transiently induced by ABA. A time course analysis of *ERF1* expression in response to ABA treatment was conducted. Wild-type seedlings were grown on MS medium vertically for 7 days, and then the seedlings were transferred to MS liquid medium without (control) or with different concentrations of ABA for the indicated times before RNA extraction from roots. The *ERF1* transcript level was analyzed by qRT–PCR analysis. Values are the mean ± SD (n = 3 experiments).

(B) Expression pattern of *ERF1* in response to ABA during LR development. Seven-day-old *ERF1pro::GUS* transgenic plants were treated with 0, 50, 100, or 200 nM ABA for 3 hours and stained for 6 hours before photographs were taken. Scale bars, 50 µm.

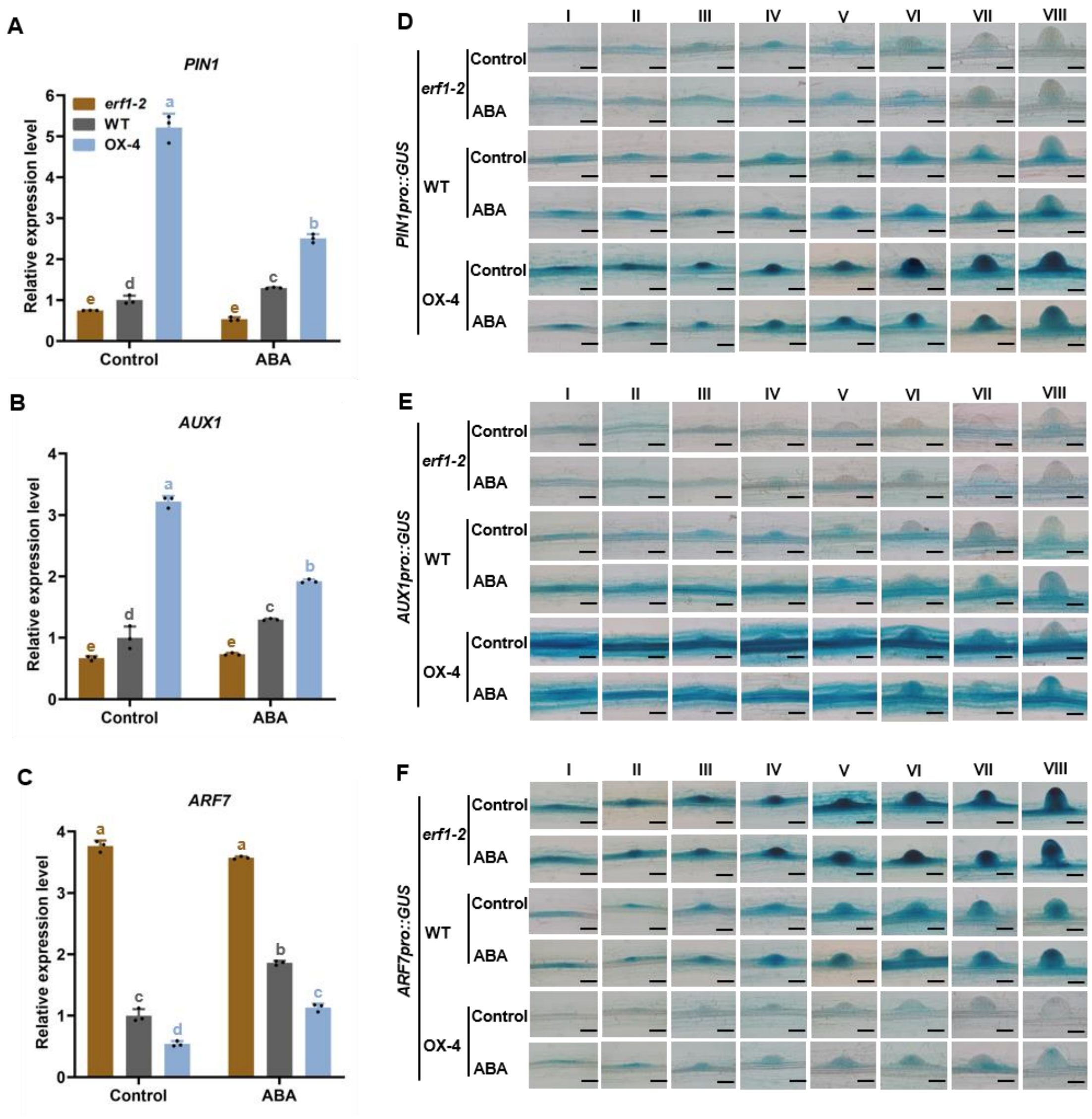


**Figure S2. The expression levels of the ERF1-targeted genes in response to ABA. Related to Figure 2.**

(A-C) Seeds of *erf1-2*, wild type, and OX-4 lines were germinated on MS medium for 7 days and transferred to MS medium with or without 200 nM ABA for 2 days. The roots from at least 40 seedlings for each line were detached, and RNA was isolated. The transcript levels of genes related to ERF1-targeted were checked by qRT–PCR analysis. The relative expression level of genes in ABA-treated samples was normalized to the WT under normal conditions. Values are the mean ± SD (n = 3 experiments). Different letters indicate significant differences by one-way ANOVA ($P < 0.05$).

(D-F) The expression of *PIN1pro::GUS*, *AUX1pro::GUS* and *ARF7pro::GUS*. Seeds of *PIN1pro::GUS*/*AUX1pro::GUS*/*ARF7pro::GUS* in the *erf-2*, WT, and OX-4 backgrounds were germinated on MS medium for 7 days, and then the seedlings were transferred to MS liquid medium without (Control) or with 200 nM ABA and incubated for 3 hours before GUS staining for 6 hours. Photographs of LR developmental stages I-VIII were taken with microscopy. Scale bar, 50 μm.

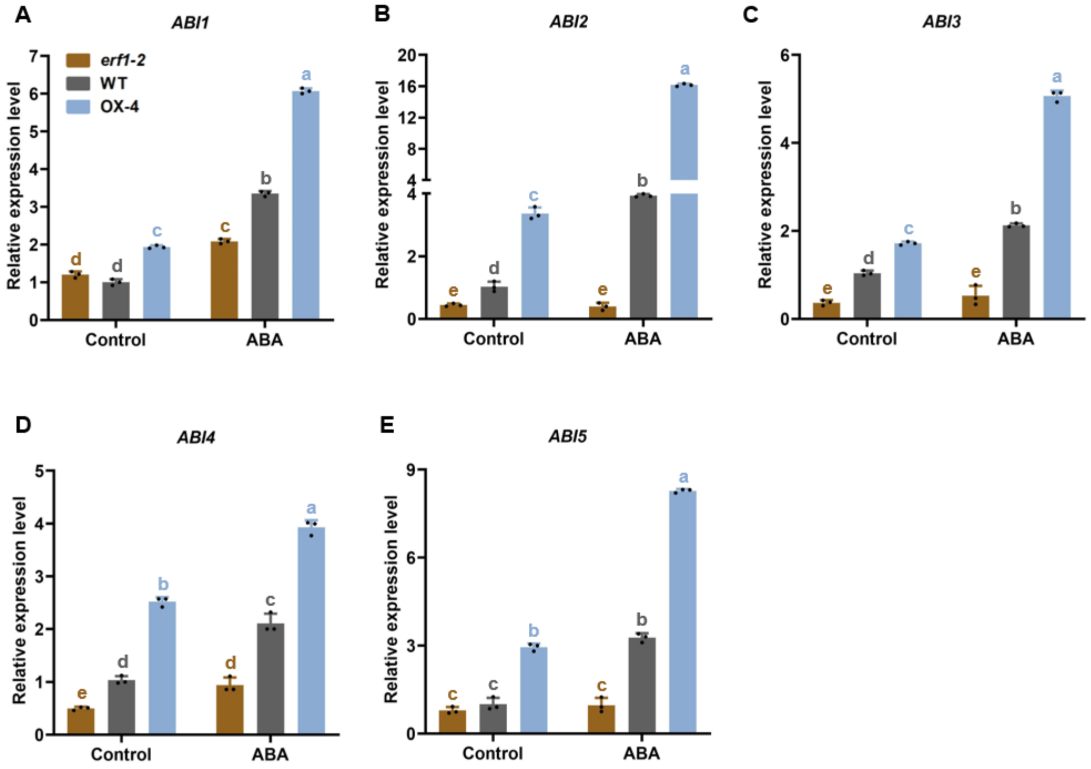


**Figure S3. The expression levels of the genes involved in ABA signaling were altered by ERF1. Related to Figure 4.**

(A-E) Seeds of *erf1-2*, WT, and OX-4 lines were germinated on MS medium for 7 days and transferred to MS medium with or without 200 nM ABA for 2 days. The roots from at least 40 seedlings for each line were detached, and RNA was isolated. The transcript levels of genes related to ABA signaling were checked by qRT–PCR analysis. The relative expression level of genes in ABA-treated samples was normalized to the WT under normal conditions. Values are the mean ± SD (n = 3 experiments). Different letters indicate significant differences by one-way ANOVA ($P < 0.05$).

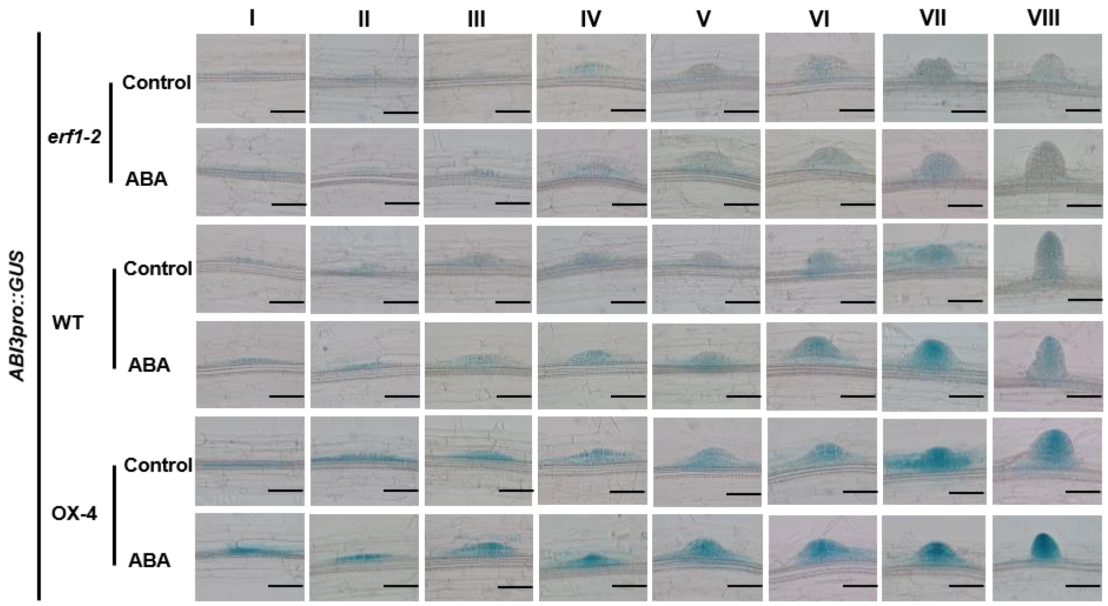


**Figure S4. The expression of *ABI3pro::GUS* was enhanced by ERF1. Related to Figure 4.** Seeds of *ABI3pro::GUS* in the *erf1-2*, WT, and OX-4 backgrounds were germinated on MS medium for 7 days, and then the seedlings were transferred to MS liquid medium without (control) or with 200 nM ABA and incubated for 3 hours before GUS staining for 6 hours. Photographs of LR developmental stages I-VIII were taken with microscopy. Scale bar, 50 µm.

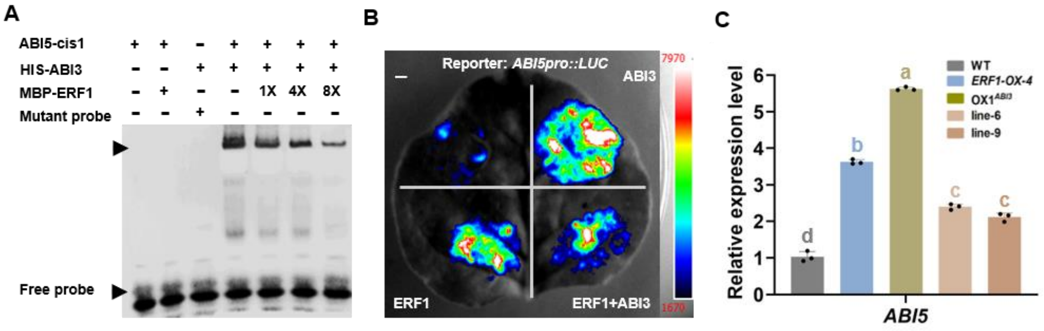


**Figure S5. ABI3-ERF1 interaction reduces the binding of ABI3 to *ABI5* promoter. Related to Figure 6.**

(A) EMSA. Biotin-labelled *ABI5-cis1* was used as probe. Unlabelled probes were used as competitors and mutated probes were used as non-competitors. + represents the presence of the component. - represents the absence of the component. As indicated, ABI3-dependent mobility shifts were detected and decreased by MBP-ERF1 protein in a dose-dependent manner. Similar results were obtained from three repeat experiments.

(B) ERF1-ABI3 interaction reduces *ABI5* trancription in transient expression assay in tobacco leaf. pRI101-*ERF1*/*ABI3* both act as effectors. pGreenII0800-*ABI5* functions as reporter. "-" indicates pRI101 empty plasmid. The different plasmid combinations were transformed into leaf cells of *N. benthamiana* leaves by agroinfiltration. The relative LUC fluorescence intensity was determined by the luciferase assay system (Tannon 5200M) at 1 days after leaf infiltration using XenolightTM d-luciferin potassium salt.

(C) The transcript levels of *ABI5*. Seeds of WT, *ERF1-OX-4*, $OX1^{ABI3}$, co-overexpression line-6 (*ERF1-OX-4* and $OX1^{ABI3}$, #6) and line-9 (*ERF1-OX-4* and $OX1^{ABI3}$, #9) were germinated on MS medium for 7 days, then RNA was isolated from roots. The relative expression level of *ABI5* was analyzed by qRT-PCR and normalized to WT. Values are mean ± SD (n=3 replicates). Different letters indicate significant difference by one-way ANOVA ($P < 0.05$).

**Table S1. Primers used in this study. Related to Figures 3, 4, 5, 6, S1, S2, S3, S4and S5.**

| Purpose | Primers | Primer sequence |
|---|---|---|
| For qRT– | *UBQ5* P1 | AGAAGATCAAGCACAAGCAT |
| PCR | *UBQ5* P2 | CAGATCAAGCTTCAACTCCT |
| | *ERF1* P1 | ATTCTTTCTCATCCTCTTCTTCT |
| | *ERF1* P2 | CGAATCTCTTATCTCCGCCG |
| | *ABI1* P1 | GTTTGGGATGTAATGACGGATG |
| | *ABI1* P2 | ACCACACTTATGTTGTCTTTGC |
| | *ABI2* P1 | CGAGATCGATGAATCAGAGTGA |
| | *ABI2* P2 | CCATCAAGCAACGAACTAGAAG |
| | *ABI3* P1 | CTGTTTCTCACCTTCAACATGG |
| | *ABI3* P2 | ATAGTTTGGAGCAGGCATGTAT |
| | *ABI4* P1 | GACTTCGTTTCATCATGAGGTG |
| | *ABI4* P2 | AGTTCAAATCCTCCATCGAACT |
| | *ABI5* P1 | CAGCTGCAGGTTCACATTCTG |
| | *ABI5* P2 | CACCCTCGCCTCCATTGTTAT |
| | *ARF7* P1 | TACCTGATGCAGCGATTGATAT |
| | *ARF7* P2 | TTCTTGTTTTAGCCGCATATCG |
| | *PIN1* P1 | TCTTCTCAAAGGCATGTATGGT |
| | *PIN1* P2 | CGAAACAATAGATCCTGCTGTG |
| | *AUX1* P1 | CCTAAGCAATTTCCTATGGCAC |
| | *AUX1* P2 | GCTCTGTATTCGACGTAGAGAA |
| For Y2H | pAD/*ERF1* P1 | GGAATTCCATATG ATGGATCCATTTTTAATTCAGTC |
| assay | pAD/*ERF1* P2 | CGGAATTCTCACCAAGTCCCACTATTTTCA |
| | pBD/*ABI3* P1 | CGGGATCCATGAAAAGCTTGCATGTGGCG |
| | pBD/*ABI3* P2 | GCGTCGACTCATTTAACAGTTTGAGAAGTTGG |
| For ChIP | *ERF1* cis1 P1 | CAGGTTTGAAGTTTAAAATAGAGCCA |
| assay | *ERF1* cis1 P2 | TATTCGAACGGATACGCAACTAAA |
| | *ERF1* control P1 | CTTCGACGAAGCAACATGGTT |
| | *ERF1* control P2 | CGGTTAGGGGATCGTTAGTGGA |
| | *ABI3* cis1 P1 | CCTTGCCTCCTTACTCACATAC |
| | *ABI3* cis1 P2 | TCTACCAACTTCATCAATTCCATC |
| | *ABI3* control P1 | AAAGAAGCTGAGACACACTTGC |
| | *ABI3* control P2 | CTATGAAATCACCTTCTTGGAG |
| For EMSA | *ARF7* cis1 +P1 | 5'biotin-CCGAGAAATTGGCGGCTCTGTCGGAG-3' |
| assay | *ARF7* cis1 -P2 | CTCCGACAGAGCCGCCAATTTCTCGG |
| | *ARF7* cis1 +P1-M | 5'biotin-CCGAGAAATTGTCTACTCTGTCGGAG-3' |
| | *ARF7* cis1 P2-M | CTCCGACAGAGTAGACAATTTCTCGG |
| | *PIN1* cis1 +P1 | 5'biotin-AGCGCACAAGGCCGCCTCTTTCACTA-3' |
| | *PIN1* cis1 -P2 | TAGTGAAAGAGGCGGCCTTGTGCGCT |
| | *PIN1* cis1 +P1-M | 5'biotin-AGCGCACAAGGTAGACTCTTTCACTA-3' |
| | *PIN1* cis1 P2-M | TAGTGAAAGACTACTCCTTGTGCGCT |
| | *AUX1* cis1 +P1 | 5'biotin-CTGCAACATTGCCGCCTTTACATAAA 3' |

| | | |
|---|---|---|
| | *AUX1* cis1 -P2 | TTTATGTAAAGGCGGCAATGTTGCAG |
| | *AUX1* cis1 P1-M | TTTATGTAAAGTAGACAATGTTGCAG |
| | *AUX1* cis1 P2-M | CTGCAACATTGTCTACTTTACATAAA |
| | *ERF1* cis1 +P1 | 5'biotin-GGACTCAGGATGCATGTGATGATGTG-3' |
| | *ERF1* cis1 -P2 | CACATCATCACATGCATCCTGAGTCC |
| | *ERF1* cis1 -P1 | GGACTCAGGATGCATGTGATGATGTG |
| | *ABI3* cis1 +P1 | 5'biotin-TTGCATGTGGCGGCCAACGCCGGA-3' |
| | *ABI3* cis1 -P2 | TCCGGCGTTGGCCGCCACATGCAA |
| | *ABI3* cis1 -P1 | TTGCATGTGGCGGCCAACGCCGGA |
| | MBP/*ABI3* P1 | AGCGGCCATGAAAAGCTTGCATGTGGCGG |
| | MBP/*ABI3* P2 | GCGTCGACTCATTTAACAGTTTGAGAAGTTG |
| | MBP/*ERF1* P1 | AGCGGCCATGGATCCATTTTTAATTCAGTC |
| | MBP/*ERF1* P2 | GCGTCGACTCACCAAGTCCCACTATTTTCA |
| For Pull-down assay | pET28a/*ABI3* P1 | TGCCGCGCGGCAGCCATATGATGAAAAGCTTGCATGTGGCG |
| | pET28a/*ABI3* P2 | CAAGCTTGTCGACGGAGCTCTCATTTAACAGTTTGAGAAGTTGG |
| | GST/*ERF1* P1 | CGGGATCCATGGATCCATTTTTAATTCAGTC |
| | GST/*ERF1* P2 | GCGTCGACTCACCAAGTCCCACTATTTTCA |
| For BiFC assay | C-pAS058/*ABI3* P1 | GCGTCGACATGAAAAGCTTGCATGTGGCG |
| | C-pAS058/*ABI3* P2 | GGACTAGTTCATTTAACAGTTTGAGAAGTT |
| | N-pAS054/*ERF1* P1 | GCGTCGACATGGATCCATTTTTAATTCAGT |
| | N-pAS054/*ERF1* P2 | GGACTAGTTCACCAAGTCCCACTATTTTCA |
| For CoIP assay | pCAM1300-HA/*ERF1* P1 | GAAGATCTATGGATCCATTTTTAATTCAGTCC |
| | pCAM1300-HA/*ERF1* P2 | GCTCTAGATCACCAAGTCCCACTATTTTCA |
| | pCAM1300-FLAG/*ABI3* P1 | GCTCTAGAATGAAAAGCTTGCATGTGGCG |
| | pCAM1300-FLAG/*ABI3* P2 | CGGGATCCTCATTTAACAGTTTGAGAAGTT |
| For tobacco LUC assay | pRI101/*ABI3* P1 | GCGTCGACATGAAAAGCTTGCATGTGGCGG |
| | pRI101/*ABI3* P2 | CGAGCTCGTCATTTAACAGTTTGAGAAGTTG |
| | pRI101/*ERF1* P1 | GCGTCGACATGGATCCATTTTTAATTCAGTCC |
| | pRI101/*ERF1* P2 | CGAGCTCGTCACCAAGTCCCACTATTTTCA |
| | pGreen0800/*ERF1* P1 | GCGTCGACATCAGTTATGCATAGTTTTGGATG |
| | pGreen0800/*ERF1* P2 | AACTGCAGGGACCCCTCTCATCGAGAAAGCA |
| | pGreen0800/*ABI3* P1 | CCCTCGAGCTCTCTCCTTTTCCTTCTGCT |
| | pGreen0800/*ABI3* P2 | GCGTCGACCTCCATAGAAGATTGAAGGGTC |
| | pGreen0800/*ARF7* P1 | GCGTCGACTTGTTCTTTGTGATCGCATATGC |
| | pGreen0800/*ARF7* P2 | AACTGCAGAATCTGAATCTGAGCTTATACAAA |
| | pGreen0800/*PIN1* P1 | CCCTCGAGCGCAACTACAACTGTAAATGAT |
| | pGreen0800/*PIN1* P2 | GCGTCGAC GTTCGCCGGAGAAGAGAGAG |

| | | |
|---|---|---|
| | pGreen0800/*AUX1* P1 | CCCTCGAGGTGGGTTGGAGTCTTGAAGAC |
| | pGreen0800/*AUX1* P2 | CCAAGCTTGTTCTTCGTTATCTTTCCCGGT |
| | pGreen0800/*ABI5* P1 | CCCTCGAG CGTGTCGAGCCTGTGAGAAGA |
| | pGreen0800/*ABI5* P2 | GCGTCGAC GATACCACCTAAACGACAATAAC |
| For expression pattern analysis | pCAMBIO/*ABI3* P1 | TGACCATGATTACGCCAAGCTTAATCAAACAATGTCATTTAGAA |
| | pCAMBIO/*ABI3* P2 | AAAACGACGGCCAGTGAATTCCCATAGAAGATTGAAGGGTCAT |
